# Automated Seizure Detection in Animal EEG Signals

**DOI:** 10.1101/2025.11.04.686618

**Authors:** Shreyan Ganguly, Zhanhong Jiang, Nyzil Massey, Nikhil Sanjay Rao, Thimmasettappa Thippeswamy, Soumik Sarkar

**Affiliations:** Iowa State University, Ames, IA, USA

## Abstract

Automated seizure detection in animal electroencephalography (EEG) is crucial for accelerating epilepsy research. While machine learning (ML) and deep learning (DL) techniques have shown promise in seizure detection for human EEG and some rodent models, their application to chemically diverse animal data remains limited. In particular, no prior work has explored deep learning approaches for EEG data where seizure/epilepsy was induced by exposure to Soman (GD), a potent agent known to generate complex and variable seizure dynamics. In this work, we evaluate deep recurrent neural networks—Gated Recurrent Units (GRU) and Long Short-Term Memory networks (LSTM)—for seizure detection in EEG signals acquired by single channel (bipotential electrodes) from the rats exposed to either kainate or GD, which later animals developed epilepsy (measured by spontaneously recurring seizures). We benchmark these models against classical baselines including Random Forest and XGBoost, using intracranial EEG data from eight animals. Our results show that GRU and LSTM substantially outperform other shallow models in both accuracy and robustness across EEG traces.

## 1 Introduction

Epilepsy is a prevalent neurological disorder characterized by recurrent, unprovoked seizures. Rodent models of epilepsy, especially chemically induced models, are extensively used to study seizure mechanisms. Electroencephalography (EEG) provides a direct and objective measure of neural activity during seizures. Traditionally, seizure detection in animal EEG recordings has relied on manual expert efforts—a process that is time-consuming as the scale of recordings grows. Recent research has begun to explore the use of machine learning (ML) and deep learning (DL) techniques for automating seizure detection [1]. These approaches offer the potential to accelerate analysis workflows, reduce human error, and improve consistency across large datasets.

While significant advances have been made in automated seizure detection for human EEG, the application of such methods to animal EEG—especially across diverse seizure-inducing conditions—remains limited. In particular, EEG datasets involving exposure to Soman (GD), a potent agent, have received little attention in the context of modern deep learning frameworks. Most existing approaches rely on handcrafted features and classical classifiers, and are often developed under homogeneous data conditions [2]. However, seizure patterns in animal models—especially those induced by agents like Soman (GD)—can exhibit considerable variability and complexity that challenge conventional methods.

Classical machine learning methods such as Support Vector Machines (SVM) [3], Random Forest [4], and XGBoost [1, 5] have been used to classify seizure activity based on handcrafted features or engineered signal features. These approaches typically rely on fixed features derived from power spectra, spike patterns, or wavelet decompositions, and have shown reasonable performance for specific seizures. Recent works have applied deep learning architectures, including fully connected neural networks [6] and convolutional neural networks (CNNs) [7], to automatically extract hierarchical features from raw or minimally processed EEG signals. While these models eliminate the need for manual feature engineering and have demonstrated improved performance, their design primarily captures local temporal dependencies.

Long Short-Term Memory (LSTM) networks have been used in related applications like vowel recognition in rodent EEG [8], suggesting their potential for temporal sequence modeling in animal data. However, their direct application to seizure detection—particularly for chemically diverse and biologically varied seizure events—remains largely under-investigated. Additionally, none of the prior works, to the best of our knowledge, have explored deep recurrent networks like LSTM or GRU for seizure detection in datasets involving GD trace. This gap in the literature motivates our work. We evaluate GRU and LSTM architectures for automated seizure detection across EEG datasets collected from multiple rodents treated with either GD or Kainate.

### Contributions

Specifically, the contributions of this paper are outlined as follows.

- We apply the popular deep recurrent models, GRU and LSTM, to detect seizures in animal EEG signals with different traces. Both GRU/LSTM empirically show more effective detection performance than the baseline ensemble learning methods. Table 1 presents qualitative comparison between our and existing works.
- Different from previous works, we establish models for diverse traces, GD and Kainate, across multiple different animals. To the best of our knowledge, our work is the first to develop deep learning-based approach for the GD trace in animal’s EEG signals.
- We use multiple datasets to empirically validate the effectiveness of trace-based deep recurrent models over different animals and traces.

**Table 1.**
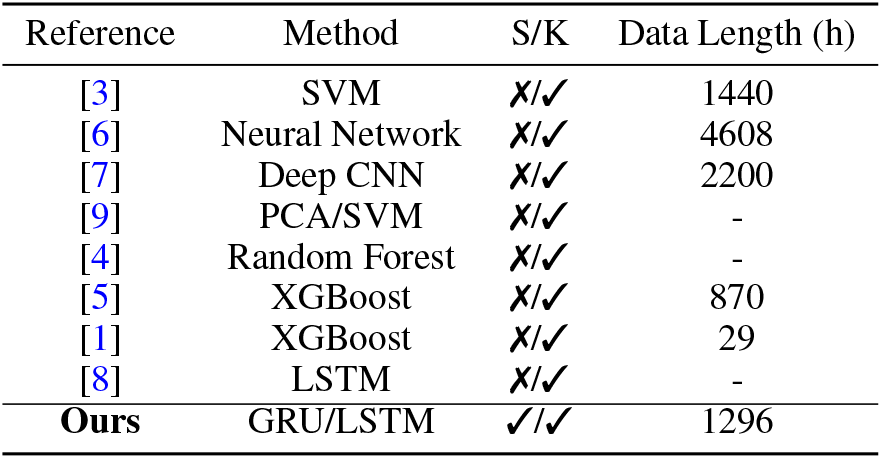
Comparison between different works. S/K: Soman (GD)/Kainate, SVM: support vector machine, CNN: convolutional neural network, PCA: principal component analysis, GRU: gated recurrent unit, LSTM: long short-term memory. Note also that we focus only on the seizure detection for rats/mice.

| Reference | Method | S/K | Data Length (h) |
| --- | --- | --- | --- |
| [3] | SVM | X/✓ | 1440 |
| [6] | Neural Network | X/✓ | 4608 |
| [7] | Deep CNN | X/✓ | 2200 |
| [9] | PCA/SVM | X/✓ | - |
| [4] | Random Forest | X/✓ | - |
| [5] | XGBoost | X/✓ | 870 |
| [1] | XGBoost | X/✓ | 29 |
| [8] | LSTM | X/✓ | - |
| **Ours** | GRU/LSTM | ✓/✓ | 1296 |

## 2 Materials and Method

The established pipeline in our work for detecting seizures in EEG signals is presented in Figure 1, which primarily involves data acquisition and processing, data segments, model training, and seizure detection. In the following, we describe each key component with more details.

**Figure 1.**
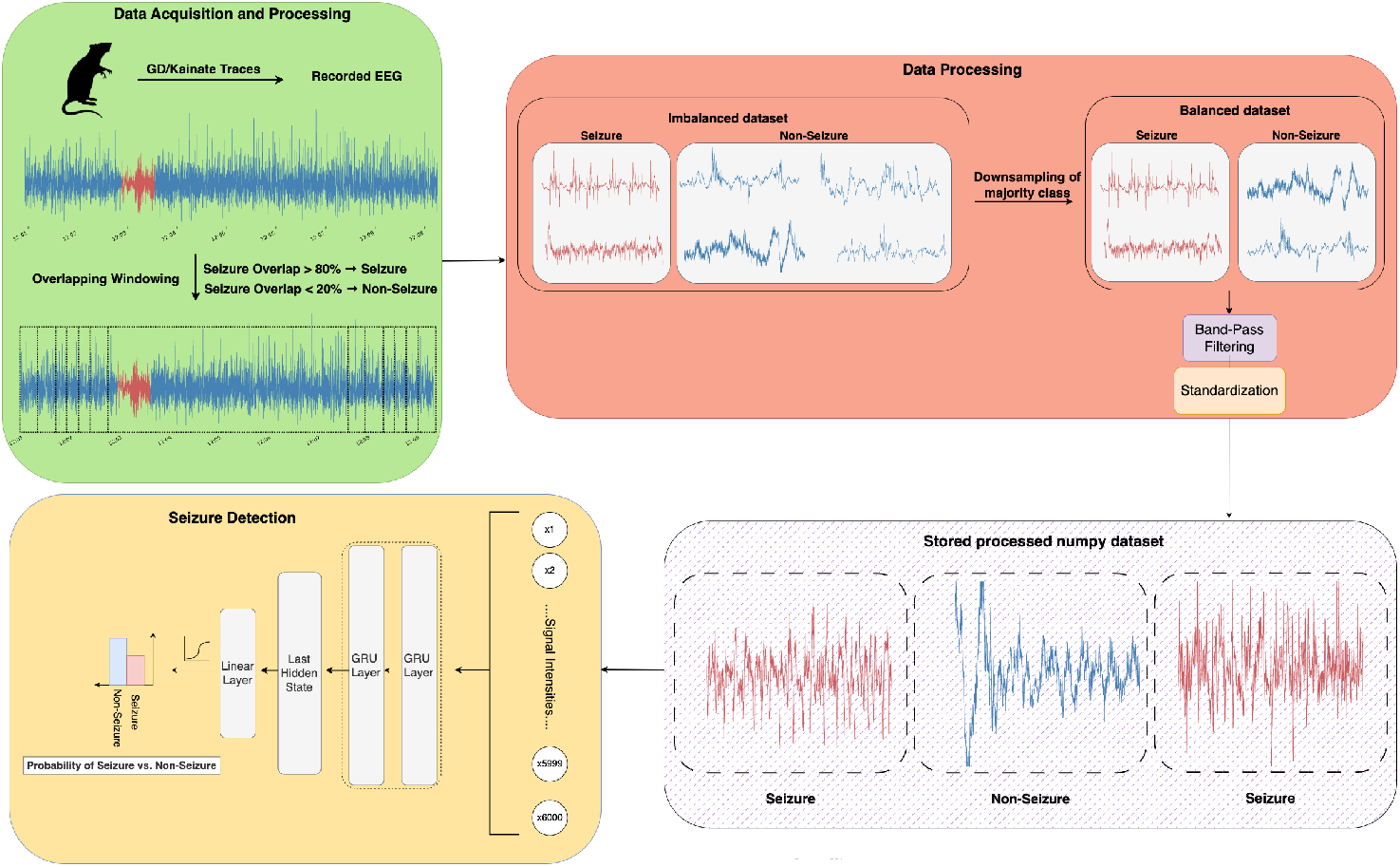
EEG Seizure detection pipeline with GRU as the example deep recurrent network.

### 2.1 Data Collection

#### Animals and Seizure Inducing Traces

The automated seizure analysis involved observing electroencephalogram (EEG) signals from eight Sprague Dawley rats (SAS400) sourced from Charles River (USA). The rats were subjected to two types of seizure inducing chemical, namely, Kainate or soman (GD). Of the eight animals, two were exposed to kainate, and six to GD. We randomly chose the EEG traces after the animals developed spontaneously recurring seizures. The total EEG duration considered in this study included 3,437,620 seconds from GD model, and 1,226,820 seconds from those treated with kainate model. A total of 4,664,440 seconds of EEG data was used in the analysis. EEG signals were recorded using intracranial single-channel electrodes implanted on the dura mater over the parietal cortex of each rat. Electrode implantation was performed under anesthesia by drilling burr holes in the parietal bones following surgical exposure via a midline scalp incision. Electrodes were secured with dental cement, and EEG signals were digitized at a sampling rate of 1000 Hz.

#### Manual Seizure Annotation

All EEG recordings were monitored and manually annotated by experts using the Ponemah Software (DSI, USA). The experts were blinded to the treatment group to remove bias. EEG seizure was defined as an episode of rhythmic spiking activity that exceeded baseline amplitude and/or frequency parameters, characterized by high-frequency and/or high-amplitude epileptiform spikes with a duration of greater than six seconds. Baseline activity was first established from seizure-free and epileptiform-free EEG segments. Seizure detection was further validated by synchronized video analysis to exclude false positives due to artifacts such as grooming, movement, or electrical noise. A total of 42,906 seconds of EEG data were annotated as seizures that included several seizure episodes with varying duration of time lapse between each episode. Of these, 35,577 seconds were observed in kainate model, and 7,329 seconds were observed in GD model (see Table 2).

**Table 2.** Overview of rat EEG recordings used in the study, including total recording duration, seizure duration, number of seizure segments in processed dataset, and inducing trace.

| Animal ID | Total Duration (s) | Seizure Duration (s) | Number of Seizure Segments (6-seconds) | Trace |
| --- | --- | --- | --- | --- |
| RN-199-23 | 240,240 | 9,377.00 | 9208 | Soman (GD) |
| RT21-21 | 431,290 | 12,480.00 | 10118 | Soman (GD) |
| RN197-23 | 343,450 | 5,779.43 | 5488 | Soman (GD) |
| RN219-23 | 931,270 | 2,337.29 | 2215 | Soman (GD) |
| RN223-23 | 645,870 | 3,042.93 | 2893 | Soman (GD) |
| RN227-23 | 845,500 | 2,558.88 | 2374 | Soman (GD) |
| RT5-24 | 362,180 | 874.00 | 844 | Kainate |
| RT2-21 | 864,640 | 6,455.00 | 2342 | Kainate |

### 2.2 Data Processing

EEG recordings, discretely spanning several days, were exported into European Data Format (.edf) files and loaded using the MNE-Python library for further processing. To generate individual units for binary seizure classification, the EEG signals were segmented into overlapping windows of six seconds, with a stride of one second. We selected six-second window length for segmentation, as it yielded the lowest false positive rate—non-seizure segments incorrectly labeled as seizures—compared to other window lengths tested up to 12 seconds. We used the same sampling rate (1000 Hz) as the original EEG recording, resulting in 6000 time points per segment. Segment labels were derived based on their temporal overlap with expert-annotated seizure intervals. Specifically, a segment was labeled as a seizure if at least 80% of its duration overlapped with a seizure annotation. Conversely, if less than 20% of a segment overlapped with seizure-labeled periods, it was labeled as a non-seizure segment. Segments falling within the intermediate overlap range (between 20% and 80%) were discarded to avoid ambiguity and reduce label noise. The total number of seizure segments obtained per animal, following this procedure, is summarized in Table 2.

Due to the sparse occurrence of seizure activity, the number of non-seizure segments was significantly higher than seizure segments. To mitigate class imbalance while preserving seizure samples, we randomly sampled non-seizure segments at a ratio of 3:1 compared to seizure segments, forming a balanced dataset. We empirically determined that a 3:1 ratio provided optimal performance across models while preserving enough class distinction during training. Each segment *X* ∈ ℝ^1*×*6000^, where 1 is the number of EEG channels and 6000 is the number of time points in the window, was standardized across the time dimension using z-score normalization:

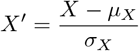

where *µ*_*X*_ ∈ ℝ^1*×*1^ and *σ*_*X*_ ∈ ℝ^1*×*1^ denote the mean and standard deviation of the segment, computed over the time axis.

### 2.3 Classification Models

To perform our analysis, we used two shallow machine learning models and two deep learning models. Among the Machine Learning models, we used ensemble learning models - Random Forest [10], and XGBoost [11]. Within the deep learning models, we used Gated Recurrent Unit (GRU) [12] and Long Short-Term Memory Network (LSTM) [13]. Random Forest and XGBoost represent the popular yet competitive models to deep learning models with significantly fewer model parameters and have been adopted in numerous applications [14]. They combine the predictions of multiple individual models (e.g., decision trees) to achieve a stronger overall prediction. For GRU and LSTM, they are all widespread deep recurrent models for modeling sequential data in either regression or classification tasks. GRU is generally faster to train and less complex than LSTM and has empirically shown more effective performance [15]. However, LSTM is more efficient in modeling long-range dependencies within the data. Surprisingly, we found that the applications of such recurrent models particularly to animals’ EEG signal seizure detection remains under-explored, remarkably motivating our study. Hyperparameters have a significant impact on deep learning model performance, which has motivated the careful tuning when designing the optimal models. In this study, we manually tuned the hyperparameters to select the nearly optimal set for different models, though an automated hyperparameter tuning method may likely be beneficial for these models. We refer interested readers to Appendix A.1 for the model description and schematic diagrams of GRU and LSTM.

#### Random Forest

Random Forest is an ensemble learning method that constructs a large number of decision trees during training and outputs the mode of their prediction for classification tasks. Each tree is trained on a bootstrapped sample of the training data and consequently, feature selection is randomized at each split. In our implementation, we used Random Forest Classifier from Scikit-Learn [16] with 100 estimator trees, and unconstrained maximum tree depth.

#### XGBoost

Extreme Gradient Boosting (XGBoost) is a gradient-boosted decision tree algorithm that builds tree sequentially, where each new tree attempts to correct errors made by previous ones. It uses second-order derivatives to improve convergence and supports regularization improving generalization.

#### Gated Recurrent Unit

Gated Recurrent Units (GRUs) are a type of recurrent neural network (RNN) that are specifically designed to handle sequential data by maintaining a hidden state over time. GRUs use gating mechanisms (reset and update gates) to control the flow of information and alleviate issues such as vanishing gradients that are common in vanilla RNNs. In our implementation, we used a two-layer GRU network with a hidden size of 64.

#### Long Short-Term Memory

Long Short-Term Memory (LSTM) networks is another variant of RNN that incorporates three gating mechanisms: input, forget, and output gates. These gates allow the LSTM to selectively retain or discard information over longer time spans, addressing the long-term dependency problem more effectively than standard RNN or GRU. In our implementation, our LSTM model consisted of two recurrent layers with a hidden size of 128.

### 2.4 Training Process

We trained models to evaluate performance at two levels of generalization: (i) within-animal, where training and evaluation data originate from the same animal, and (ii) within-trace, where data is grouped by chemical trace (Kainate or GD). For both settings, the processed EEG data was grouped accordingly, and models were trained and evaluated under these constraints to assess generalization across biological and experimental contexts. We randomly split the balanced dataset into 70% training and 30% testing subsets, ensuring class balance within each set.

The deep-learning classifiers (GRU and LSTM) were trained for 120 epochs. The models were trained using the Adam optimizer with a learning rate of 0.001 and binary cross-entropy loss. A batch size of 32 was used across all experiments. During training, model checkpoints were saved every 20 epochs, and the checkpoint achieving the highest evaluation metric on the full unbalanced segment set was selected as the corresponding final model. All experiments were run with a fixed random seed (42) to ensure reproducibility of training and evaluation. The same training and evaluation protocol was applied to the shallow classifiers (Random Forest and XGBoost). Both Random Forest and XGBoost classifiers were trained using default hyperparameters from the respective Scikit-Learn and XGBoost libraries.

### 2.5 Evaluation Metrics

Model performance was evaluated on both the created balanced test set and the full unbalanced segment set. The balanced test set enables comparison under controlled class distribution, while the unbalanced evaluation set reflects the true deployment scenario, where seizure events are sparse. Specifically, we used the following evaluation metrics to assess the performance of every saved checkpoints: Accuracy, ROC AUC (Area Under the Receiver Operating Characteristic Curve), Precision, and Recall.

#### Accuracy

Accuracy measures the overall correctness of the classifier and is defined as the proportion of correctly predicted segments among all segments:

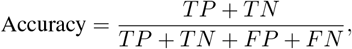

where *TP* denotes true positives (correctly predicted seizure segments), *TN* true negatives (correctly predicted non-seizure segments), *FP* false positives (non-seizure segments incorrectly predicted as seizure), and *FN* false negatives (seizure segments incorrectly predicted as non-seizure).

#### ROC AUC

The ROC AUC evaluates the trade-off between the true positive rate (sensitivity) and the false positive rate across all possible classification thresholds. It is a threshold-independent measure that summarizes model discriminability:

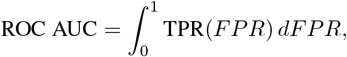

where TPR = *TP/*(*TP* + *FN*) and FPR = *FP/*(*FP* + *TN*). A value of 1.0 indicates perfect separation between classes, while 0.5 implies no discriminative power.

#### Precision

Precision quantifies how many of the segments predicted as seizure are truly seizure:

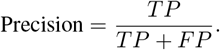

High precision indicates a low false positive rate.

#### Recall

Recall (also known as Sensitivity or True Positive Rate) measures how many of the true seizure segments were correctly identified by the model:

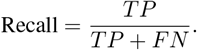

High recall is crucial for seizure detection tasks, as missing actual seizure events can have significant consequences.

## 3 Results and Discussion

We evaluated four classification models—Random Forest, XGBoost, GRU, and LSTM—for automated seizure detection across multiple experimental settings. Performance was assessed at both the individual animal level and the trace level, using the discussed classification metric. In this section, all results are reported on the full unbalanced test set, which reflects real-world seizure frequency. Evaluation on the balanced set showed similar relative performance trends across models.

### 3.1 Animal-Level Performance

Table 3 and Table 4 summarize the accuracy and AUC ROC scores achieved by different models at the individual animal level. GRU and LSTM consistently outperformed the shallow models across both metrics. On average, LSTM achieved an accuracy of 98.78%, followed by GRU with 98.62% and Random Forest of 97.86% across all animals. GRU obtained the highest accuracy in five out of eight animals, including 99.70% on RN197-23 and 99.35% on RT21-21. AUC ROC results followed a similar trend. GRU achieved an average AUC ROC of 99.40% across all animals, slightly higher than LSTM (99.27%) and significantly better than Random Forest (96.94%) and XGBoost (97.76%). The results indicate good robustness of GRU in modeling temporal patterns associated with seizures across both GD and Kainate traces.

**Table 3.** Accuracy (in %) of different models trained and evaluated at individual animal-level.

| Animal ID | Random Forest | XGBoost | GRU | LSTM |
| --- | --- | --- | --- | --- |
| RN-199-23 | 97.49 | 98.43 | **99.03** | 98.57 |
| RT21-21 | 99.50 | 99.03 | **99.35** | 99.10 |
| RN197-23 | 99.28 | 99.61 | **99.70** | 99.52 |
| RN219-23 | 97.54 | 98.16 | 98.72 | **98.84** |
| RN223-23 | 97.71 | 99.31 | 98.76 | **99.67** |
| RN227-23 | 95.63 | 98.04 | 98.41 | **98.63** |
| RT5-24 | 96.89 | **98.10** | 95.96 | 97.10 |
| RT2-21 | 98.90 | **99.25** | 99.08 | 98.84 |

**Table 4.** AUC ROC (in %) of different models trained and evaluated at individual animal-level.

| Animal ID | Random Forest | XGBoost | GRU | LSTM |
| --- | --- | --- | --- | --- |
| RN-199-23 | 94.56 | 98.37 | **99.79** | 99.55 |
| RT21-21 | 95.62 | 97.86 | **99.83** | 99.07 |
| RN197-23 | 97.74 | 99.54 | **99.96** | 99.74 |
| RN219-23 | 95.57 | 96.41 | 98.59 | **98.89** |
| RN223-23 | 96.54 | 98.43 | **98.83** | 99.67 |
| RN227-23 | 95.21 | 96.57 | 99.65 | **99.71** |
| RT5-24 | 95.18 | 95.06 | 97.25 | **98.01** |
| RT2-21 | 99.64 | 98.20 | 99.31 | **99.52** |

Figure 2a illustrates the recall achieved by each model when trained and evaluated separately for each animal. GRU consistently outperformed all other models, achieving recall more than 90% for most animals. LSTM also showed high recall, though slightly lower and less consistent. In contrast, the shallow models (Random Forest and XGBoost) exhibited more variability and generally lower recall values, especially on GD-treated animals. GRU’s high recall across all animals highlights its effectiveness in capturing seizure patterns across diverse EEG profiles. Even in animals with fewer seizure events, such as RT5-24, GRU maintained competitive sensitivity.

**Figure 2.**
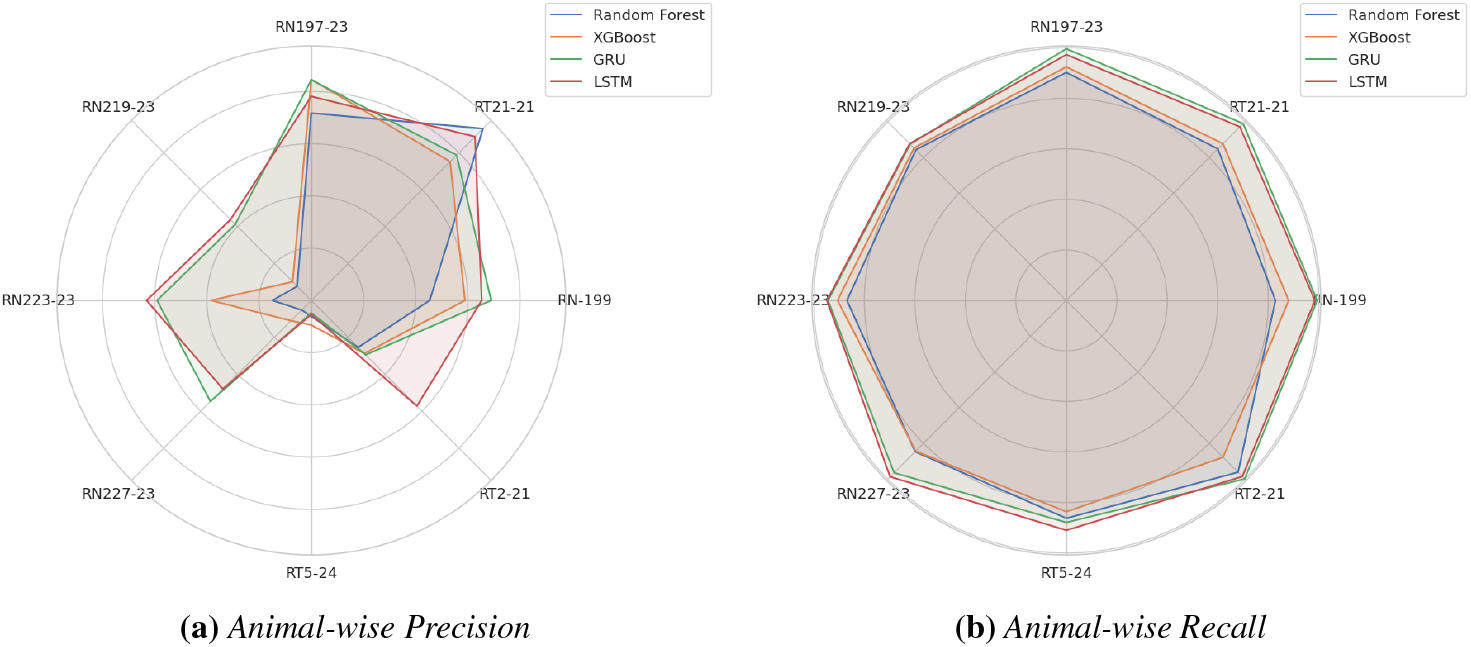
This figure compares the precision (left) and recall (right) of four classifiers (Random Forest, XGBoost, GRU, and LSTM) when trained and evaluated separately on each animal’s EEG data. GRU and LSTM models generally achieve higher recall and precision across most animals, particularly those exposed to Soman (GD), highlighting their effectiveness in modeling subject-specific seizure dynamics.

Figure 2b illustrates the precision of the same models. GRU achieved the highest precision across most animals, with values reaching 78.73% for RT21-21. LSTM performed comparably in certain cases (e.g., RT2-21), while shallow models exhibited significantly lower precision—particularly in GD-treated animals such as RN219-23 and RN227-23, where Random Forest and XGBoost dropped below 15%. While seizure detection in Kainate-treated animals (e.g., RT5-24, RT2-21) remains challenging for all models, GRU and LSTM still outperform shallow baselines. The drop in precision for these animals may stem from signal variability or limited seizure-labeled segments in the training data.

### 3.2 Trace-Level Performance

To evaluate model generalization across seizure-inducing agents, we grouped data by trace (GD vs. Kainate) and trained/evaluated models accordingly. Figure 3 summarizes recall and precision at the trace level. GRU achieved the highest recall across both traces, with values exceeding 94.1% on GD and 93.4% on Kainate. LSTM followed closely but exhibited greater variance in Kainate, consistent with its per-animal performance. Shallow models were less reliable compared to the deep networks. In terms of precision, GRU outperformed other models, achieving 48.2% and 46.57% for GD and Kainate, respectively. LSTM’s precision dropped notably on Kainate, while Random Forest and XGBoost struggled across both traces. These results confirms the benefit of recurrent models for modeling temporal EEG patterns and detecting rare seizure events.

**Figure 3.**
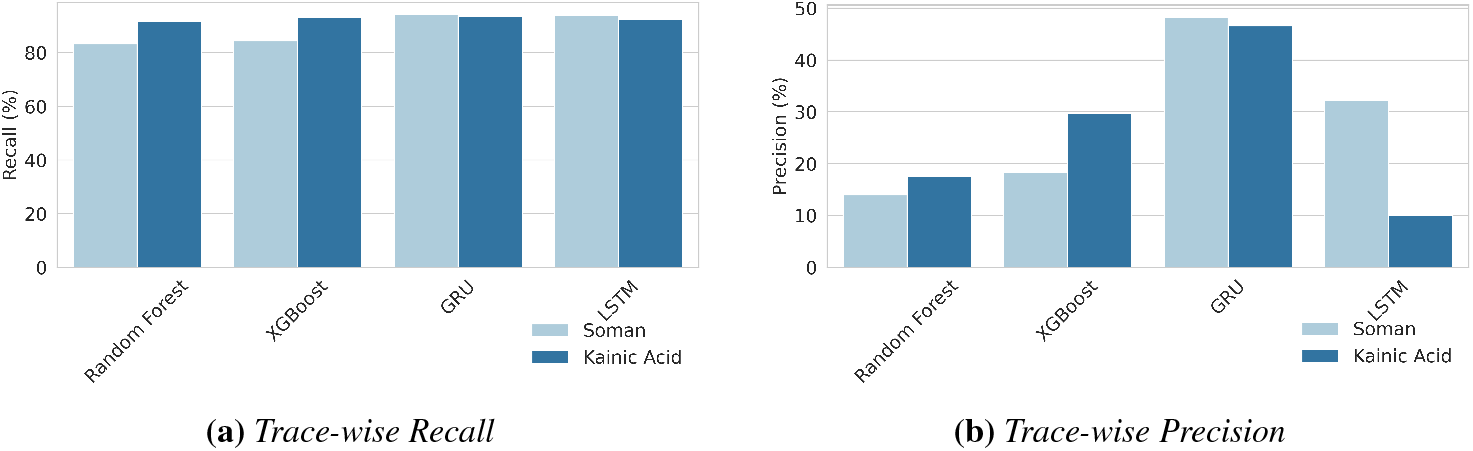
The figure shows the average precision (right) and recall (left) for all models evaluated on grouped EEG data from animals exposed to either Soman (GD) or Kainate. GRU consistently outperforms other models on both traces, indicating its robustness across chemically distinct seizure patterns.

### 3.3 Training

Figure 4 illustrates the training and validation loss curves for GRU over 120 epochs on the GD dataset. Loss decreased rapidly in the first 20 epochs and stabilized thereafter. The validation loss remained low and relatively flat, indicating strong generalization and absence of overfitting. The stability of the loss curves supports the robustness of the GRU architecture and confirms the effectiveness of our training protocols.

**Figure 4.**
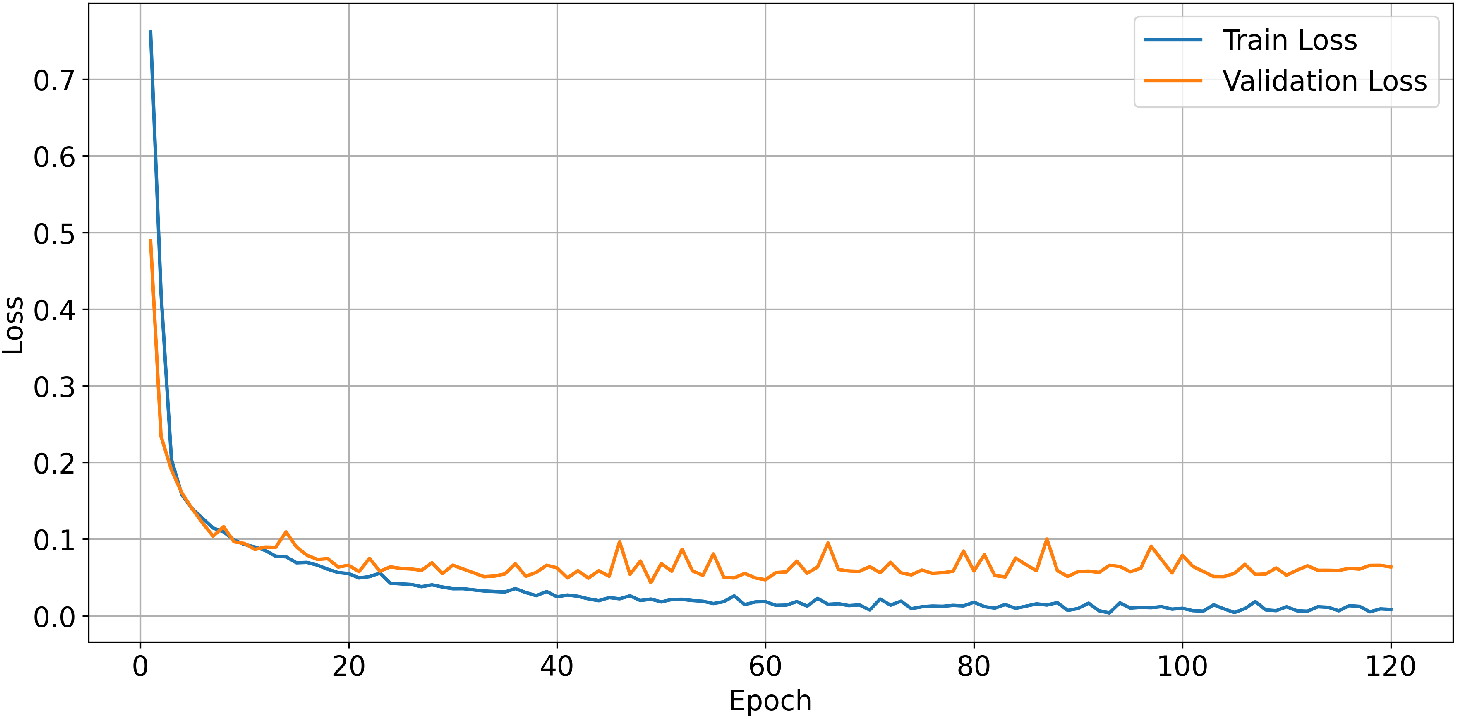
The figure shows the binary cross-entropy loss over 120 training epochs.

### 3.4 Discussion

Our findings demonstrate the strong potential of GRU-based models for automated seizure detection in animal EEG data. GRU consistently achieved higher recall and precision across individual animals and chemical traces. Its ability to model temporal dynamics using gated mechanisms enables it to effectively distinguish seizure from non-seizure segments, under biological and experimental variability. Shallow models like Random Forest and XGBoost offered computational efficiency but failed to capture the temporal features necessary for accurate seizure classification. These models may remain useful in constrained environments or when paired with domain-specific signal features, but they are clearly outperformed by recurrent deep learning approaches in raw EEG modeling.

## 4 Conclusion

We presented an automated deep learning approach for seizure detection in animal EEG signals, evaluating GRU and LSTM models across diverse chemical traces and rodent subjects. Our results show that recurrent neural networks, particularly GRUs, substantially outperform traditional ensemble classifiers in both accuracy and robustness. This performance is especially compelling for GD-induced EEG data, which presents significant variability and has not been previously studied using deep learning. Notably, the GRU model achieved consistently high recall across animals and traces—indicating a strong ability to detect seizure events when they occur. This high recall supports the reliability of the model in minimizing missed detections, a critical requirement for automated seizure analysis. These findings position deep recurrent architectures as reliable and scalable tools for seizure detection in preclinical epilepsy research. Future directions include extending the approach to non-convulsive seizure and interictal or epileptiform spikes detection, multi-channel EEG, and systematic study of model performance on inter-animal and intra-animal training and evaluation schemes.

## A Additional Technical Appendices

### A.1 LSTM and GRU

We detail the technical information about GRU and LSTM in this section for completeness.

#### LSTM

We briefly review the working mechanism of LSTM in this context. As a popular variant of recurrent neural network (RNN) in sequence modeling, LSTM, as depicted in Figure 5, successfully resolves the vanishing gradient issues in vanilla RNN. At the time step *k*, we denote by **x**_*k*_, **h**_*k*_ and **c**_*k*_ respectively the input vector, the hidden state and the cell state. For each time step, LSTM calculates the cell state **c**_*k*_, the input gate **i**_*k*_, the forget gate **f**_*k*_ and the output gate **o**_*k*_, resulting in the following set of equations.

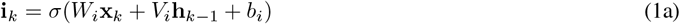

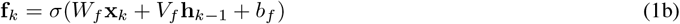

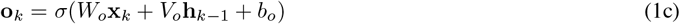

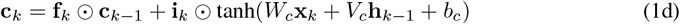

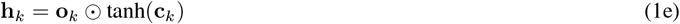

**Figure 5.**
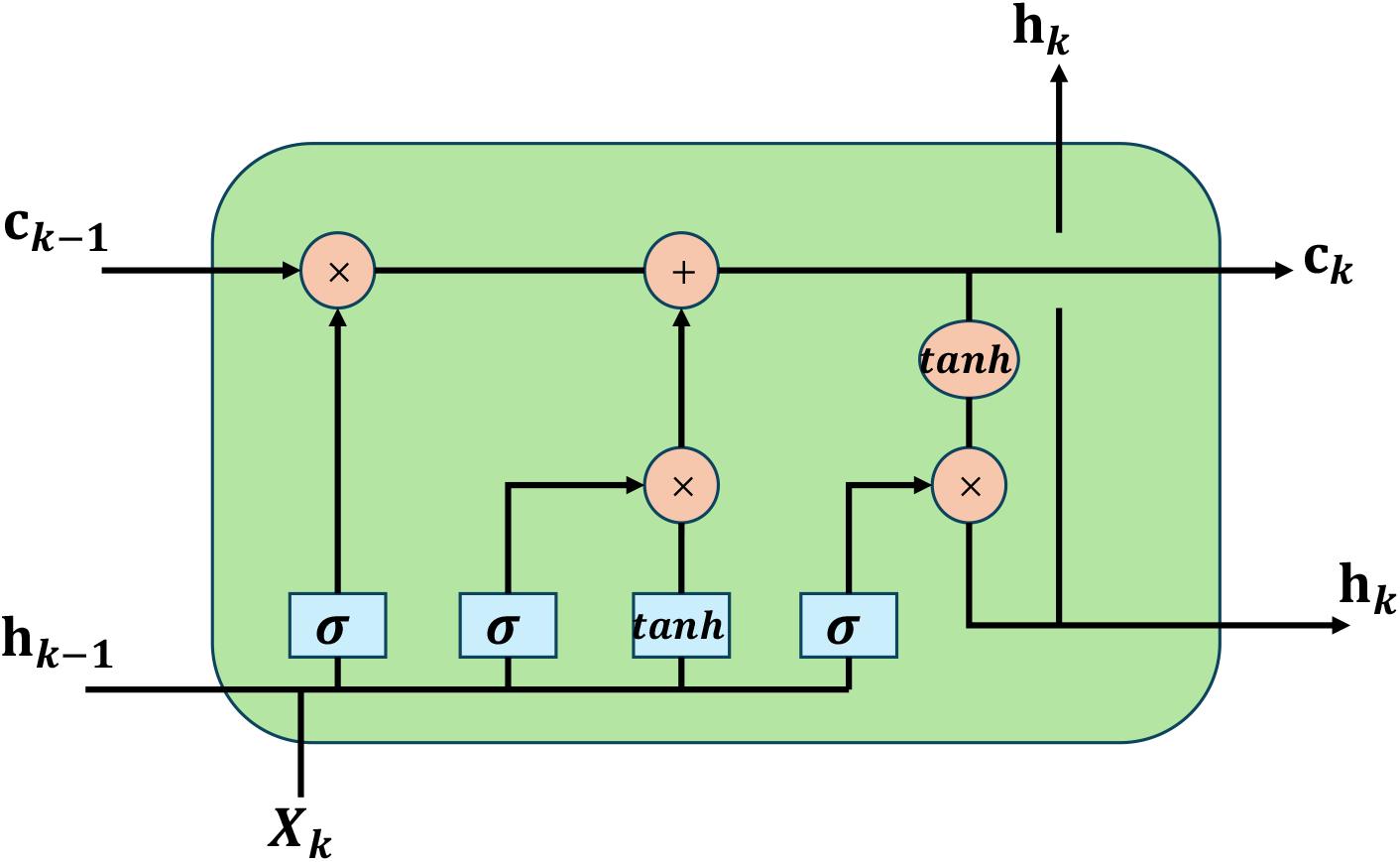
Schematic Diagram of LSTM.

**Figure 6.**
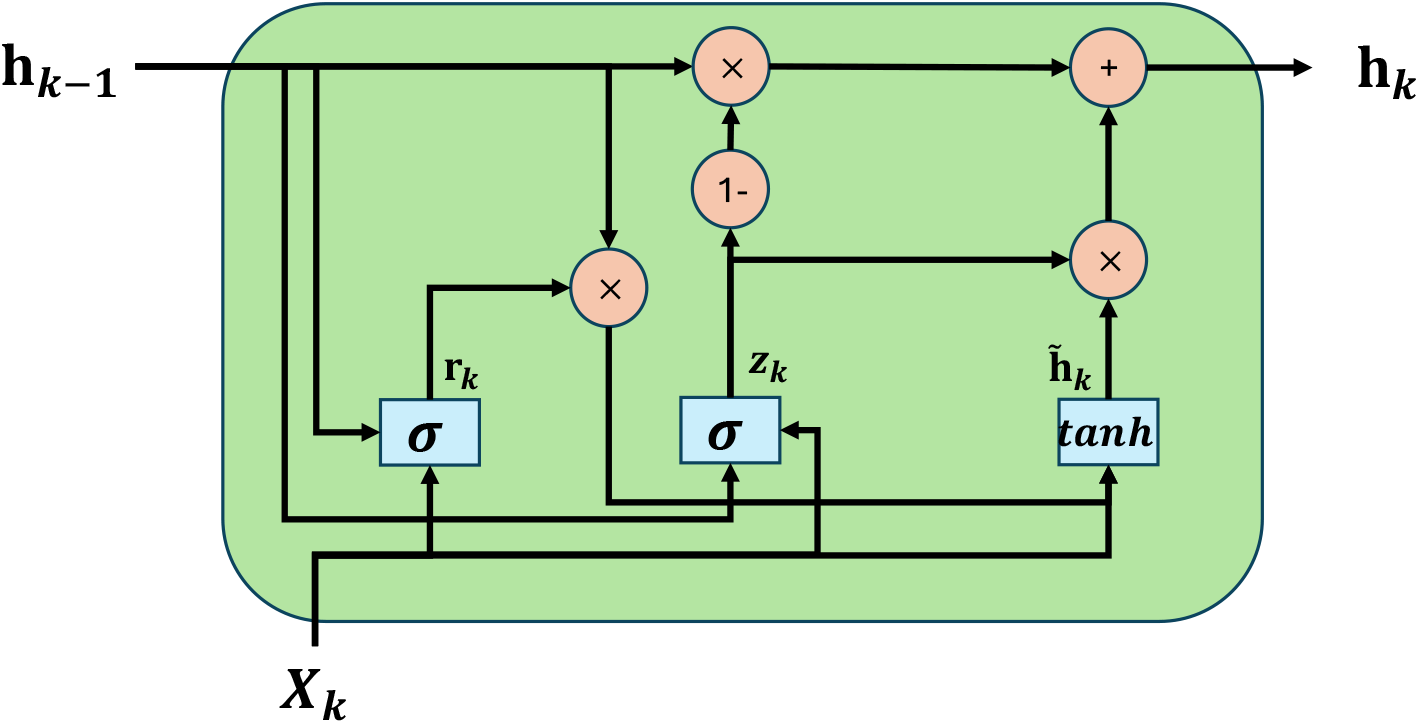
Schematic Diagram of GRU.

*W* _∗_, *V*_∗_ correspond to weight matrices regarding different gates and state. *b*_∗_ are the bias vectors. ⊙ is the element-wise product. Intuitively, LSTM effectively learns when to remember and when to forget pertinent information in a sequence, ensuring effective information processing when modeling a long-range dependencies.

#### GRU

GRU is a gating mechanism in RNNs, proposed in 2014 by Cho et al. [17], which is similar to LSTM but lacks a context vector **c** or output gate **o**. This naturally results in fewer parameters than LSTM. Specifically, the internal update law is summarized as follows:

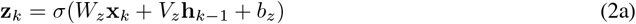

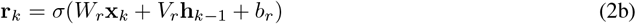

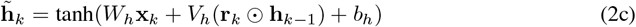

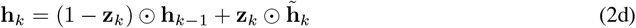

Similarly, *W* _∗_, *V*_∗_ correspond to weight matrices regarding different gates. **z** is called the update gate while **r** reset gate. All elements of these two gate vectors are between 0 and 1. Due to the simpler mechanism of GRU, it may be less effective at capturing very long-term dependencies compared to LSTM. However, in practice, it has shown competitive performance to LSTM in multiple areas, such as timeseries analysis, speech signal modeling and natural language processing [18–20].

